# Reconstructing the backbone of the Saccharomycotina yeast phylogeny using genome-scale data

**DOI:** 10.1101/070235

**Authors:** Xing-Xing Shen, Xiaofan Zhou, Jacek Kominek, Cletus P. Kurtzman, Chris Todd Hittinger, Antonis Rokas

## Abstract

Understanding the phylogenetic relationships among the yeasts of the subphylum Saccharomycotina is a prerequisite for understanding the evolution of their metabolisms and ecological lifestyles. In the last two decades, the use of rDNA and multi-locus data sets has greatly advanced our understanding of the yeast phylogeny, but many deep relationships remain unsupported. In contrast, phylogenomic analyses have involved relatively few taxa and lineages that were often selected with limited considerations for covering the breadth of yeast biodiversity. Here we used genome sequence data from 86 publicly available yeast genomes representing 9 of the 11 major lineages and 10 non-yeast fungal outgroups to generate a 1,233-gene, 96-taxon data matrix. Species phylogenies reconstructed using two different methods (concatenation and coalescence) and two data matrices (amino acids or the first two codon positions) yielded identical and highly supported relationships between the 9 major lineages. Aside from the lineage comprised by the family Pichiaceae, all other lineages were monophyletic. Most interrelationships among yeast species were robust across the two methods and data matrices. However, 8 of the 93 internodes conflicted between analyses or data sets, including the placements of: the clade defined by species that have reassigned the CUG codon to encode serine, instead of leucine; the clade defined by a whole genome duplication; and of *Ascoidea rubescens.* These phylogenomic analyses provide a robust roadmap for future comparative work across the yeast subphylum in the disciplines of taxonomy, molecular genetics, evolutionary biology, ecology, and biotechnology. To further this end, we have also provided a BLAST server to query the 86 Saccharomycotina genomes, which can be found at http://y1000plus.org/blast.

## Introduction

Molecular phylogenetic analyses show that the fungal phylum Ascomycota is comprised of three monophyletic subphyla that share a common ancestor ~500 million years ago (Kurtzman and Robnett 1994; Sugiyama *et al.* 2006; Taylor and Berbee 2006; James *et al.* 2006; Liu *et al.* 2009): the Saccharomycotina (syn. Hemiascomycota; e.g. *Saccharomyces, Pichia, Candida*), the Pezizomycotina (syn. Euascomycota; e.g. *Aspergillus, Neurospora*) and the Taphrinomycotina (syn. Archaeascomycota; e.g. *Schizosaccharomyces, Pneumocystis*).

Yeasts of the fungal subphylum Saccharomycotina exhibit remarkably diverse heterotrophic metabolisms, which have enabled them to successfully partition nutrients and ecosystems and inhabit every continent and every major aquatic and terrestrial biome (Hittinger *et al.* 2015). While yeast species were historically identified by metabolic differences, recent studies have shown that many of these classic characters are subject to rampant homoplasy, convergence, and parallelism (Hittinger *et al.* 2004; Hall and Dietrich 2007; Wenger *et al.* 2010; Slot and Rokas 2010; Lin and Li 2011; Wolfe *et al.* 2015). Despite the considerable progress in classifying yeasts using multi-locus DNA sequence data, critical gaps remain (Kurtzman and Robnett 1998, 2003, 2007, 2013; Nguyen *et al.* 2006; Kurtzman *et al.* 2008, 2011; Kurtzman and Suzuki 2010); many genera are paraphyletic or polyphyletic, while circumscriptions at or above the family level are often poorly supported (Hittinger *et al.* 2015).

In recent years, phylogenomic analyses based on data matrices comprised of hundreds to thousands of genes from dozens of taxa have provided unprecedented resolution to several, diverse branches of the tree of life (Song *et al.* 2012; Salichos and Rokas 2013; Liang *et al.* 2013; Xi *et al.* 2014; Wickett *et al.* 2014; Whelan *et al.* 2015). Although the genomes of several dozen yeast species are currently available (Hittinger *et al.* 2015), published phylogenomic studies contain at most 25 yeast genomes (Rokas *et al.* 2003; Fitzpatrick *et al.* 2006; Liu *et al.* 2009; Medina *et al.* 2011; Salichos and Rokas 2013; Marcet-Houben and Gabaldón 2015; Riley *et al.* 2016).

A robustly resolved backbone yeast phylogeny will be of great benefit, not only to the study of yeast biodiversity, but also to diagnosticians seeking to identify and treat yeast infections, to biotechnologists harnessing yeast metabolism to develop advanced biofuels, and to biologists designing computational and functional experiments. Toward that end, here we have used genome sequence data from 86 publicly available yeast genomes representing 9 of the 11 major lineages and 10 non-yeast fungal outgroups to reconstruct the backbone of the Saccharomycotina yeast phylogeny.

## Materials and Methods

### Data acquisition

The workflow used to assemble the data sets for the inference of the backbone phylogeny of Saccharomycotina yeasts is described in Figure 1. To assemble a dataset with the greatest possible taxonomic sampling as of January 11th, 2016, we first collected all Saccharomycotina yeast species whose genomes were available (Hittinger *et al.* 2015). We then excluded 4 publicly available genomes, namely, *Blastobotrys attinorum, Blastobotrys petasosporus, Cephaloascus albidus*, and *Cephaloascus fragrans*, which had been released under embargo and lacked a citable publication. In addition, we excluded the genomes of known hybrid species, such as *Pichia farinosa* (Louis *et al.* 2012), *Saccharomyces cerevisiae × Saccharomyces eubayanus* syn. *Saccharomyces pastorianus* (Libkind *et al.* 2011; Gibson and Liti 2015), and the wine yeast VIN7 *(Saccharomyces cerevisiae × Saccharomyces kudriavzevii)* (Borneman *et al.* 2012). For species with multiple isolates sequenced, we only included the genome of the isolate with the highest number of the “complete” genes (see below). These criteria resulted in the inclusion of genomes from 86 yeast species representing 9 of 11 major lineages of the subphylum Saccharomycotina (Hittinger *et al.* 2015). Finally, we used the genomes of 10 non-yeast fungi that are representatives of the phylum Ascomycota as outgroups. Detailed information of the nomenclature, taxonomy, and source of the 96 genomes in our study is provided in Table S1.

**Figure 1:**
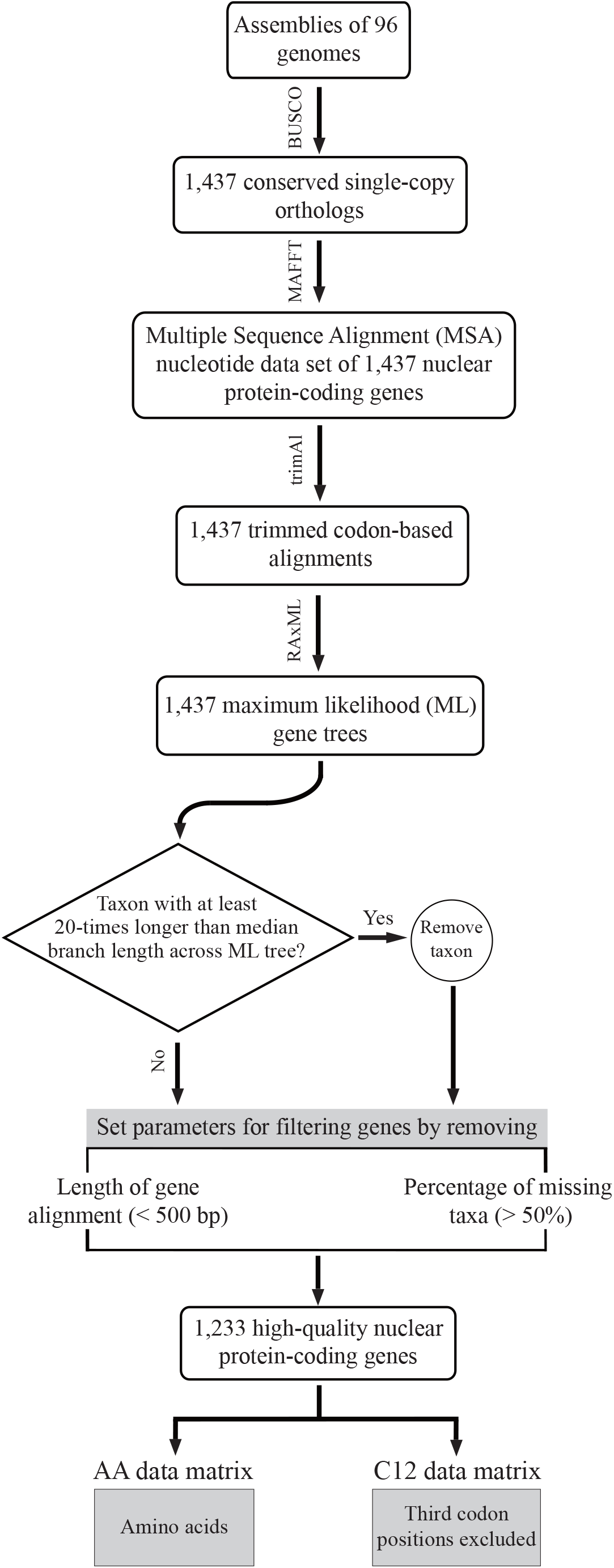
Workflow illustrating the steps involved in the construction of the two phylogenomic data matrices used in this study.

### A custom BLAST database for the genomes of the 86 yeast species

To further facilitate the use of these 86 Saccharomycotina genomes by the broader research community, we set up a custom local BLAST database using Sequenceserver, version 1.0.8 (Priyam *et al.* 2015). The database is free and publicly available through http://y1000plus.org/blast.

### Assessment of genome assemblies and ortholog identification

Assessment of the 96 selected genome assemblies was performed using the BUSCO software, version 1.1b (Simão *et al.* 2015). Each individual genome was examined for the copy number of 1,438 pre-selected genes (hereafter, referred to as BUSCO genes) that are single-copy in at least 90% of the 125 reference fungal genomes in the OrthoDB Version 7 database (www.orthodb.org) (Waterhouse *et al.* 2013). Briefly, for each BUSCO gene, a consensus protein sequence was generated from the hidden Markov model (HMM) alignment profile of the orthologous protein sequences among the 125 reference genomes using the HMMER software, version 3 (Eddy 2011). This consensus protein sequence was then used as query in a tBLASTn (Altschul *et al.* 1990; Camacho *et al.* 2009) search against each genome to identify up to three putative genomic regions, and the gene structure of each putative genomic region was predicted by AUGUSTUS (Stanke and Waack 2003). Next, the sequences of these predicted genes were aligned to the HMM alignment profile of the BUSCO gene, and the ones with alignment bit-score higher than a pre-set cutoff (90% of the lowest bit-score among the 125 reference genomes) were kept. If no predicted gene from a particular genome was retained after the last step, the gene was classified as “missing” from that genome. If one or more predicted genes from a genome were retained, these were further examined for their aligned sequence length in the HMM-profile alignment; predicted genes whose aligned sequence lengths were shorter than 95% of the aligned sequence lengths of genes in the 125 reference genomes were classified as “fragmented”. The remaining predicted genes were classified as “complete” if only one predicted gene was present in a genome, and “duplicated” if two or more “complete” predicted genes were present in a genome. Only the sequences of single-copy “complete” genes without any in-frame stop-codon(s) were used to construct ortholog groups across the 96 genomes. We excluded the orthologous group constructed from BUSCO gene “BUSCOfEOG7MH16D” from our subsequent analyses because sequences of this gene consistently failed to be predicted by AUGUSTUS across the 96 genomes.

### Sequence alignment, alignment trimming, and removal of spurious sequences and low-quality genes

For each ortholog group, we first translated nucleotide sequences into amino acid sequences using a custom Perl script, taking into account the differential meaning of the CUG codon in the CUG-Ser clade of yeasts whose CUG codon encodes serine, instead of leucine (Dujon 2010; Mühlhausen and Kollmar 2014; Hittinger *et al.* 2015; Riley *et al.* 2016). Next, we aligned the amino acid sequences using the E-INS-i strategy as implemented by the program MAFFT, version 7.215 (Katoh and Standley 2013), with the default gap opening penalty (--op = 1.53). We then used a custom Perl script to map the nucleotide sequences on the amino acid alignment and to generate the codon-based nucleotide alignment. Regions of ambiguous alignment in codon-based nucleotide alignments were trimmed using the trimAl software, version 1.4 (Capella-Gutierrez *et al.* 2009) with the ‘gappyout’ option on; otherwise, default settings were assumed. Finally, the trimmed codon-based alignments were translated into trimmed amino acid alignments.

To minimize the inclusion of potentially spurious or paralogous sequences, the maximum likelihood (ML) phylogram for the trimmed codon sequence alignment of each ortholog group was inferred under an unpartitioned ‘GTR (Tavaré 1986) + GAMMA (Yang 1994, 1996)’ model as implemented in RAxML, version 8.2.3 (Stamatakis 2014). Sequences whose terminal branch (leaf) lengths were at least 20-times longer than the median of all terminal branch lengths across the maximum likelihood (ML) phylogram for a given orthologous group were excluded. In total, 49 sequences from 42 ortholog groups were removed. The resulting gene alignments were further filtered by length of trimmed gene alignment (alignments that were less than 500 base pairs in length were removed from downstream analyses) and taxon number (alignments with less than 50% gene occupancy, i.e., that contained fewer than 48 taxa, were removed from downstream analyses).

The remaining 1,233 ortholog groups were used to generate two data matrices: A) a C12 data matrix that included only the first and second codon positions of every gene (third codon positions were excluded because they showed much higher variation in GC content than the first and second codon positions; Figure S1); and B) an AA data matrix that included the amino acid sequences of every gene.

### Phylogenetic analysis

Phylogenetic analysis was performed separately for the AA and C12 data matrices using the ML optimality criterion (Felsenstein 1981), under two different approaches: concatenation (Huelsenbeck *et al.* 1996; Rokas *et al.* 2003; Philippe *et al.* 2005) and coalescence (Edwards 2009). To infer the concatenation phylogeny for the AA data matrix, we used an unpartitioned ‘PROTGAMMALG’ model of amino acid substitution, as 681 out of 1,233 genes favored ‘LG (Le and Gascuel 2008) + Gamma (Yang 1994, 1996)’ for rate heterogeneity among sites as best-fitting model (Figure S2). To infer the concatenation phylogeny for the C12 data matrix, we used an unpartitioned ‘GTR (Tavaré 1986) + GAMMA (Yang 1994, 1996)’ model of nucleotide substitution. In both cases, phylogenetic reconstruction was performed using 5 randomized maximum parsimony trees and 5 random trees as starting trees in RAxML (Stamatakis 2014). Branch support for each internode was evaluated with 100 rapid bootstrapping replicates (Stamatakis *et al.* 2008). Finally, we also used a gene-based partition scheme (1,233 partitions) to separately conduct ML tree search for the AA and C12 data matrices, in which parameters of evolutionary model (amino acid; see below; DNA, GTR+G) were separately estimated for each orthologous group (-q option) in RAxML. As the ML trees produced by the gene-based partition scheme on both data matrices were topologically identical to the ML trees produced by the unpartitioned scheme (results are deposited on the figshare repository), and the computational resources required for partitioned analyses are much greater, bootstrap support and phylogenetic signal analyses were performed using only the unpartitioned scheme.

For the coalescence-based analyses of the AA data matrix, the best-fitting model of amino acid evolution for each orthologous group was selected using the Bayesian Information Criterion (BIC) (Schwarz 1978), as implemented in ProtTest 3.4 (Darriba *et al.* 2011). For the C12 data matrix, the GTR + GAMMA model was used to accommodate nucleotide substitution and rate heterogeneity among sites. In both cases, we inferred the best-scoring unrooted ML gene tree for every ortholog group by running 10 separate ML searches using RAxML. Branch support for each internode was evaluated with 100 rapid bootstrapping replicates (Stamatakis *et al.* 2008). Individually estimated ML gene trees were used as input to estimate the coalescent-based phylogenies for the AA and C12 data matrices using the ASTRAL software, version 4.7.7 (Mirarab *et al.* 2014). The robustness of these phylogenies was evaluated by the multi-locus bootstrap approach (Seo 2008) with 100 replicates, each of which consisted of individual gene trees each selected randomly from the set of 100 rapid bootstrapping trees available for each gene. Finally, the set of the 1,233 ML gene trees was used to calculate (partial) internode certainty (IC) values (Salichos *et al.* 2014; Kobert *et al.* 2016), as implemented in RAxML, version 8.2.3.

### Selecting subsets of genes with strong phylogenetic signal

Selecting strongly supported genes has been empirically shown to reduce incongruence among gene trees in phylogenetic analyses (Salichos and Rokas 2013; Wang *et al.* 2015). To quantify support for individual gene trees, we used two common phylogenetic measures on the AA data matrix: 1) the average bootstrap support (ABS), which was calculated using a custom Perl script, and corresponds to the average of bootstrap support values across the ML tree of a given gene; 2) the relative tree certainty (RTC), which was calculated in RAxML, and corresponds to the average of all internode certainty (IC) values across the ML tree of a given gene (Salichos *et al.* 2014; Kobert *et al.* 2016). IC values on each ML tree were calculated by examining the bipartitions present in the topologies generated by the 100 rapid bootstrap replicates.

Four subsets of 1,233 genes in the AA data matrix were constructed based on the ABS and RTC measures, respectively: the first two subsets included the 616 genes (top 50%) having the highest ABS or RTC values, respectively; the remaining two subsets included the 308 genes (top 25%) having the highest ABS or RTC values, respectively. Since the subsets constructed using the ABS and RTC phylogenetic measures showed no significant differences in the sets of genes included (Figure S3), we used the two subsets (top 50% and top 25%) based on ABS for subsequent analyses. For each subset, the concatenation phylogeny and the coalescent-based phylogeny, as well as their clade support values (bootstrap support and IC values), were separately estimated using RAxML and ASTRAL by following the procedures described above.

### Data availability

All data matrices and their resulting phylogenies have been deposited on the figshare repository at DOI: 10.6084/m9.figshare.3370675.

## Results and Discussion

### Genome completeness

Contig or scaffold N50 (i.e., the contig or scaffold size above which 50% of the total length of the sequence assembly can be found) is by far the most widely used measure to assess the quality of a genome assembly (Yandell and Ence 2012). The higher N50 is, the better the assembly generally is. Nonetheless, this value does not assess assembly completeness in terms of gene content. Thus, we assessed completeness for each of 96 fungal genomes using the BUSCO set of 1,438 single-copy, conserved genes among 125 fungi (Simão *et al.* 2015). We found that the percentage of “complete” BUSCO genes among the 96 genomes ranged from 65.2 % to 98.5% of 1,438 fungal BUSCO genes, with the average being 94.2% (Figure 2, Table S2 for detailed values). Only 10 of the 96 fungal genomes had less than 90% of 1,438 fungal BUSCO genes, with the assemblies of *Hanseniaspora valbyensis* (65.2%) and *Hanseniaspora uvarum* (67.5%) having the lowest coverage, with ~ 300 BUSCO genes either missing or fragmented in each of the two genomes. Since the ancestral internode length leading to the two *Hanseniaspora* genomes was among the longest in the yeast phylogeny (Figure 3), their low coverage may be a consequence of this accelerated evolutionary rate.

**Figure 2:**
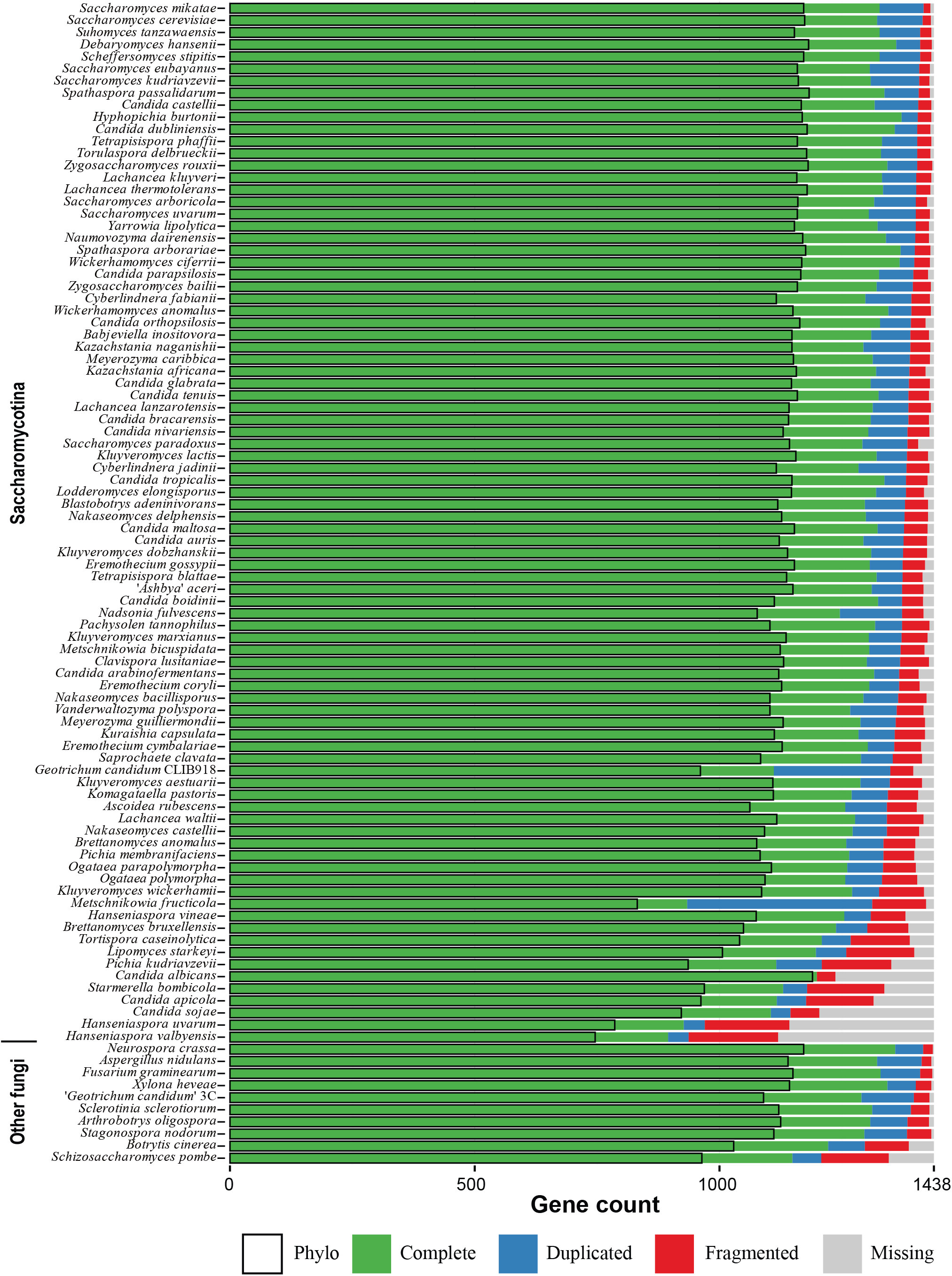
Genomic quality assessment of the 86 yeast and 10 outgroup fungal genomes used in this study. The bar plot next to each species indicates the fractions of BUSCO genes that are present or missing in each genome. “Complete”: fraction of single-copy, full-length genes; “Duplicated”: fraction of multiple-copy, full-length genes; “Fragmented”: fraction of genes with a partial sequence; “Missing”: fraction of genes not found in the genome; “Phylo”: fraction of single-copy “Complete” genes used to construct the phylogenomic data matrices. Yeast species are arranged along the Y-axis in ascending order of the total number of “Fragmented” and “Missing” genes. The exact value of quality assessment of each species can be found in Table S2.

**Figure 3:**
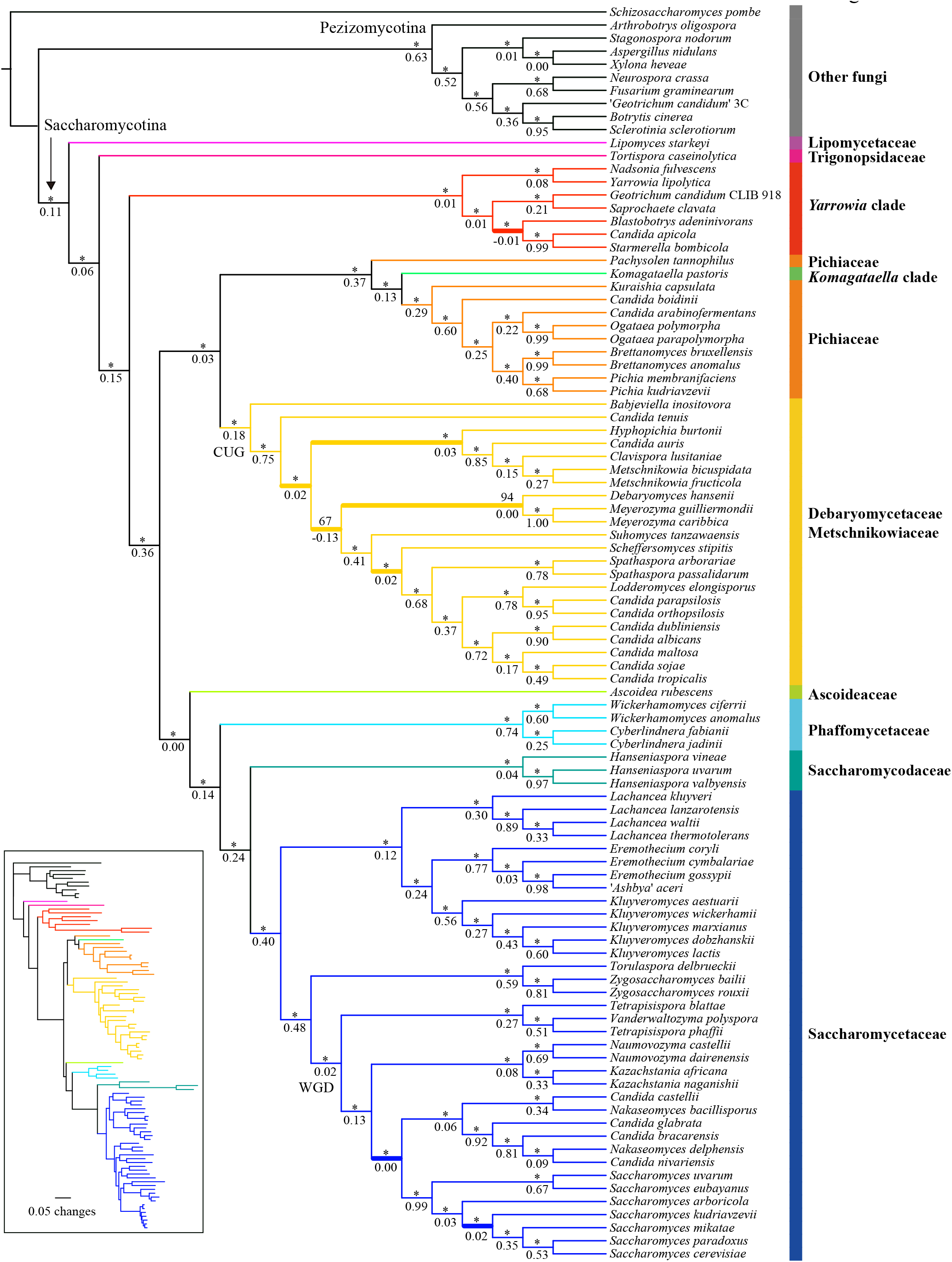
The phylogenetic relationships of Saccharomycotina yeasts inferred from the concatenation-based analysis of a 1,233 single-copy BUSCO gene amino acid (AA) data matrix. The ML phylogeny was reconstructed based on the concatenation amino acid data matrix (609,899 sites) under an unpartitioned LG + GAMMA substitution model using RAxML version 8.2.3 (Stamatakis 2014). Branch support values near internodes are indicated as bootstrap support value (above) and internode certainty (below), respectively. * indicates bootstrap support values greater than or equal to 95%. Thicker branches show conflicts between concatenation-based phylogeny (Figure 3) and coalescence-based phylogeny (Figure 4). Note, branch lengths on the ML tree are given in the ***inset*** at the bottom left.

**Figure 4:**
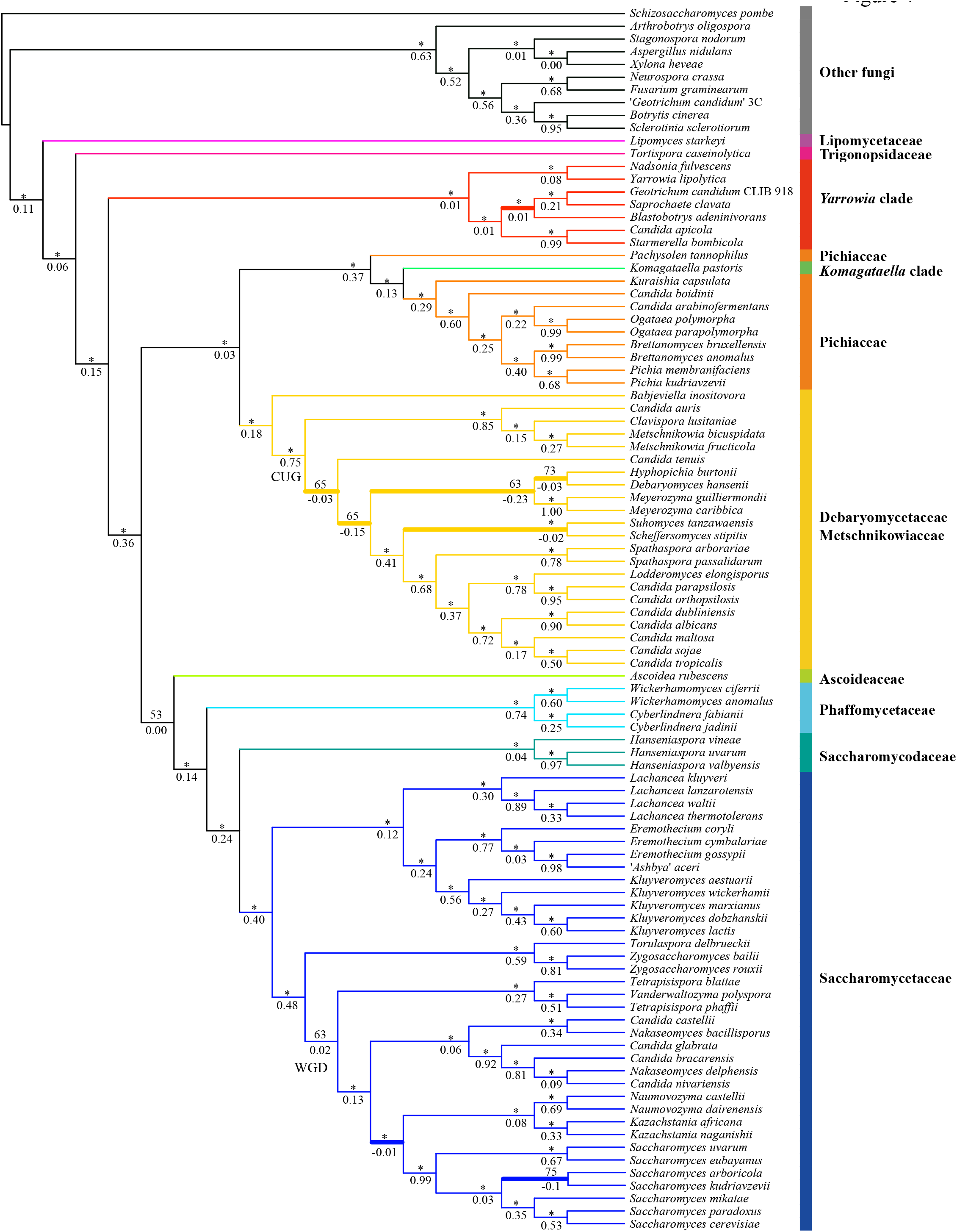
The phylogenetic relationships of Saccharomycotina yeasts inferred from the coalescence-based analysis of a 1,233 single-copy BUSCO gene amino acid (AA) data matrix. The coalescence-based phylogeny estimation was conducted using ASTRAL version 4.7.7 (Mirarab *et al.* 2014). Branch support values near internodes are indicated as bootstrap support value (above) and internode certainty (below), respectively. * indicates bootstrap support values greater than or equal to 95%. Thicker branches show conflicts between coalescence-based phylogeny (Figure 4) and concatenation-based phylogeny (Figure 3).

### Data matrix completeness

Following orthology identification, alignment trimming, and removal of spurious sequences and low-quality genes, we retained 1,233 of the original 1,438 orthologous groups from the 96 genomes (see Materials and Methods). None of the genomes of the 96 species had all 1,233 orthologous groups, but 88 had more than 1,000 orthologous groups each (Table S2, Figure 4). The percentage of gene occupancy in the remaining 8 species ranged from 60.5% (746 / 1,233) to 78.6% (969 / 1,233). Two recent studies showed that nonrandom bias in missing data might be potentially problematic for phylogenetic inference (Xi *et al.* 2016; Hosner *et al.* 2016), particularly for coalescence-based phylogenetic inference (Hovmöller *et al.* 2013). Interestingly, the placements for the 8 low gene-coverage species in our study were robust in both concatenation and coalescent analyses, as well as in the two data matrices and subsampling analyses (see results below), suggesting that the impact of missing data in this study is negligible.

Among the 1,233 orthologous groups, the percentage of sequence occupancy ranged from 50% to 100%, with an average value of 90%; 1,084 orthologous groups displayed at least 80% sequence occupancy, and 24 contained sequences from all 96 species (Figure S4, Table S3). In addition, gene alignment lengths ranged from 501 to 14,562 base pairs (bp), with an average length of 1,484 bp (Figure S5, Table S3). The AA and C12 data matrices contained a total of 609,899 and 1,219,798 sites, respectively.

### A genome-wide yeast phylogeny

All phylogenetic analyses consistently separated the 10 non-yeast outgroup taxa (nine Pezizomycotina and one Taphrinomycotina species) from the Saccharomycotina yeasts (Figures 3 and 4, Figures S6 and S7). Surprisingly, the genomes of two yeast isolates purported to be from the same species *(Geotrichum candidum* isolates CLIB 918 and 3C) were placed into two different clades; *Geotrichum candidum* CLIB 918 (Morel *et al.* 2015) was nested within Saccharomycotina yeasts, whereas '*Geotrichum candidum*’ 3C (Polev *et al.* 2014) was nested within the Pezizomycotina outgroup. Given its phylogenetic placement, we infer that the genome sequence of the isolate '*Geotrichum candidum*’ 3C represents a misidentified Pezizomycotina species.

Most internodes in the concatenation phylogenies inferred from the AA and C12 data matrices received high bootstrap support values that were equal or greater than 95% (AA, 91 out of 93 internodes; C12, 88 out of 93 internodes) (Figure 3, Figure S6). Similar to the results of the concatenation approach, the two coalescence-based phylogenies were also robustly supported, with 86 / 93 internodes (AA data matrix) and 86 / 93 internodes (C12 data matrix) showing 95% or greater bootstrap support (Figure 4, Figure S7). There were 5 conflicting internodes between the AA and C12 concatenation-based phylogenies and 2 conflicting internodes between the AA and C12 coalescence-based phylogenies. In addition, comparison of the concatenation-based phylogeny to the coalescence-based phylogeny showed 8 topological differences in the phylogenies inferred using the AA data matrix and 5 in the phylogenies inferred using the C12 data matrix. These topological differences are discussed below.

### Stable and conflicted internodes

Of the 9 major Saccharomycotina lineages, 8 were monophyletic in all analyses; the only exception was the family Pichiaceae. This lineage was paraphyletic because *Komagataella pastoris*, which belongs to the *Komagataella* clade, groups within it (Figure 3). Overall, the family Lipomycetaceae was resolved as the earliest-branching lineage of Saccharomycotina yeasts, followed by the family Trigonopsidaceae and the *Yarrowia* clades (Figure 3). A clade consisting of the family Pichiaceae, the CUG-Ser clade, and the *Komagataella* clade was well supported. The family represented by the most genome sequences, the Saccharomycetaceae, was recovered as the sister group to the family Saccharomycodaceae. This Saccharomycetaceae/Saccharomycodaceae clade was sister to the Phaffomycetaceae. These relationships were mostly recovered in two multigene studies (Kurtzman and Robnett 2013; Mühlhausen and Kollmar 2014) and one phylogenomic study based on 25 yeast genomes (Riley *et al.* 2016), and they are broadly consistent with the most recent views of the yeast phylogeny (Hittinger *et al.* 2015). Finally, 8 of the 9 major Saccharomycotina lineages were robustly and strongly supported in both concatenation- and coalescence-based analyses. The only exception was the family Ascoideaceae, whose support values under concatenation analyses were high (AA: BS = 95%; C12: BS = 97%) but were much lower under coalescence-based analyses (AA: BS = 53%; C12 = 45%). This finding is consistent with the instability in the placement of *Ascoidea rubescens* inferred from multiple analyses of different data matrices in the original genome study (Riley *et al.* 2016). Thus, although this family was consistently recovered as the closest relative of the Saccharomycetaceae/Saccharomycodaceae/Phaffomycetaceae clade in our analyses (Figures 3 and 4, Figures S6 and S7), its current placement in the yeast phylogeny should be considered tenuous and unresolved.

To quantify incongruence between the 1,233 orthologous groups across the 93 internodes of the yeast phylogeny, we used the 1,233 individual ML gene trees to calculate internode certainty (IC) values (Salichos and Rokas 2013; Salichos *et al.* 2014; Kobert *et al.* 2016). Our results showed that most of the internodes in concatenation- and coalescence-based phylogenies inferred from AA and C12 data matrices had IC values greater than 0 (Figures 3 and 4, Figures S6 and S7), suggesting that those relationships were recovered by the majority of 1,233 genes. For the aforementioned internodes that were incongruent between approaches (concatenation versus coalescence) or data matrices (AA versus C12), their IC values were often less than 0 (Figures 3 and 4, Figures S6 and S7). These conflicting internodes occurred within the WGD/allopolyploidization clade (Wolfe and Shields 1997; Marcet-Houben and Gabaldón 2015) in the family Saccharomycetaceae (3 topological differences), the CUG-Ser clade (4 topological differences), and the *Yarrowia* clade (1 topological difference).

Within the WGD clade, the concatenation phylogenies identified the *Nakaseomyces* clade as the sister group to the genus *Saccharomyces* (Figure 3, Figure S6), a relationship supported by several rare genomic changes (Scannell *et al.* 2006), whereas the phylogenies inferred from the coalescence-based approach strongly supported the *Kazachstania/Naumovozyma* clade as the sister group to the genus *Saccharomyces* (Figure 4, Figure S7), a relationship supported by phylogenomic (Salichos and Rokas 2013) and multigene (Mühlhausen and Kollmar 2014) analyses. The monophyly of yeasts for the WGD clade was recovered by the concatenation- and coalescence-based phylogenies on the AA data matrix, as well as by the concatenation-based phylogeny on the C12 data matrix, consistent with recent studies (Dujon 2010; Salichos and Rokas 2013; Wolfe *et al.* 2015). In contrast, the coalescence-based phylogeny on the C12 data matrix weakly supported a paraphyly of the WGD clade, in which a ZT clade composed of the genus *Zygosaccharomyces* and the genus *Torulaspora* nested within the WGD clade in agreement with the results from multigene studies (Kurtzman and Robnett 2013; Mühlhausen and Kollmar 2014). Interestingly, a recent examination of 516 widespread orthologs from 25 yeast genomes inferred that the WGD event was the result of an allopolyploidization between the KLE clade (the genus *Kluyveromyces*, the genus *Lachancea*, and the genus *Eremothecium)* and the ZT clade (Marcet-Houben and Gabaldón 2015), providing a potential explanation for the observed instability of the WGD clade. A sister relationship between *Saccharomyces arboricola* and *Saccharomyces kudriavzevii* was only recovered by the coalescence-based phylogeny inferred from the AA data matrix (Figure 4); this relationship contrasts with all other phylogenies inferred in this study, as well as those obtained in most published studies (Scannell *et al.* 2011; Liti *et al.* 2013; Hittinger 2013; Mühlhausen and Kollmar 2014; Hittinger *et al.* 2015).

Within the CUG-Ser clade, *Suhomyces (Candida) tanzawaensis* was strongly recovered as sister to *Scheffersomyces stipitis* by both methods on the C12 data matrix, as well as by the coalescence-based phylogeny on the AA data matrix. In contrast, the concatenation-based phylogeny using the AA data matrix strongly supported a sister relationship between *Su. tanzawaensis* and a clade composed of *Sc. stipitis*, the genus *Spathaspora*, and some of yeasts within the CUG-Ser clade, in agreement with the results of the original study describing the *Su. tanzawaensis* genome (Riley *et al.* 2016). The closest relatives of the genus *Meyerozyma* were either *Debaryomyces hansenii, Candida tenuis*, or a clade containing *Debaryomyces hansenii* and *Hyphopichia burtonii*, depending on the analysis and data matrix considered. Finally, within the *Yarrowia* clade, the concatenation phylogenies inferred from AA and C12 data matrices placed the *Candida apicola/Starmerella bombicola* clade as the sister group to *Blastobotrys (Arxula) adeninivorans* (Figure 3, Figure S6), whereas the coalescence-based phylogenies inferred from AA and C12 data matrices supported *Blastobotrys (Arxula) adeninivorans* sister to the *Saprochaete clavata/Geotrichum candidum* isolate CLIB 918 clade (Figure 4, Figure S7). Finally, the concatenation- and coalescence-based phylogenies inferred from AA and C12 data matrices consistently recovered a sister group of *Nadsonia fulvescens* and *Yarrowia lipolytica*, but its resolution in coalescence-based phylogenies inferred from C12 data matrices received weak median bootstrap support (BS = 65%). This group was not recovered in the previous multigene study of these taxa (Kurtzman and Robnett 2013).

### Selecting genes with strong phylogenetic signal reduces incongruence

To examine whether the use of genes with strong phylogenetic signal could reduce incongruence among individual gene trees (Salichos and Rokas 2013; Wang *et al.* 2015), we constructed two AA data matrices containing the 308 (top 25%) or 616 (top 50%) ortholog groups showing the highest average bootstrap values in their bootstrap consensus gene trees and reconstructed their phylogenies by concatenation and coalescence. The IC values of most internodes in both of the concatenation-based (ML) phylogeny (all orthologous groups, average IC value = 0.41; top 50%, average IC value = 0.55; top 25%, average IC value = 0.64) and the coalescence-based phylogeny (all orthologous groups, average IC value = 0.40; top 50%, average IC value = 0.54; top 25%, average IC value = 0.65) increased (Figure 5), suggesting that the use of genes with strong phylogenetic signal decreased the amount of incongruence in the yeast phylogeny.

**Figure 5:**
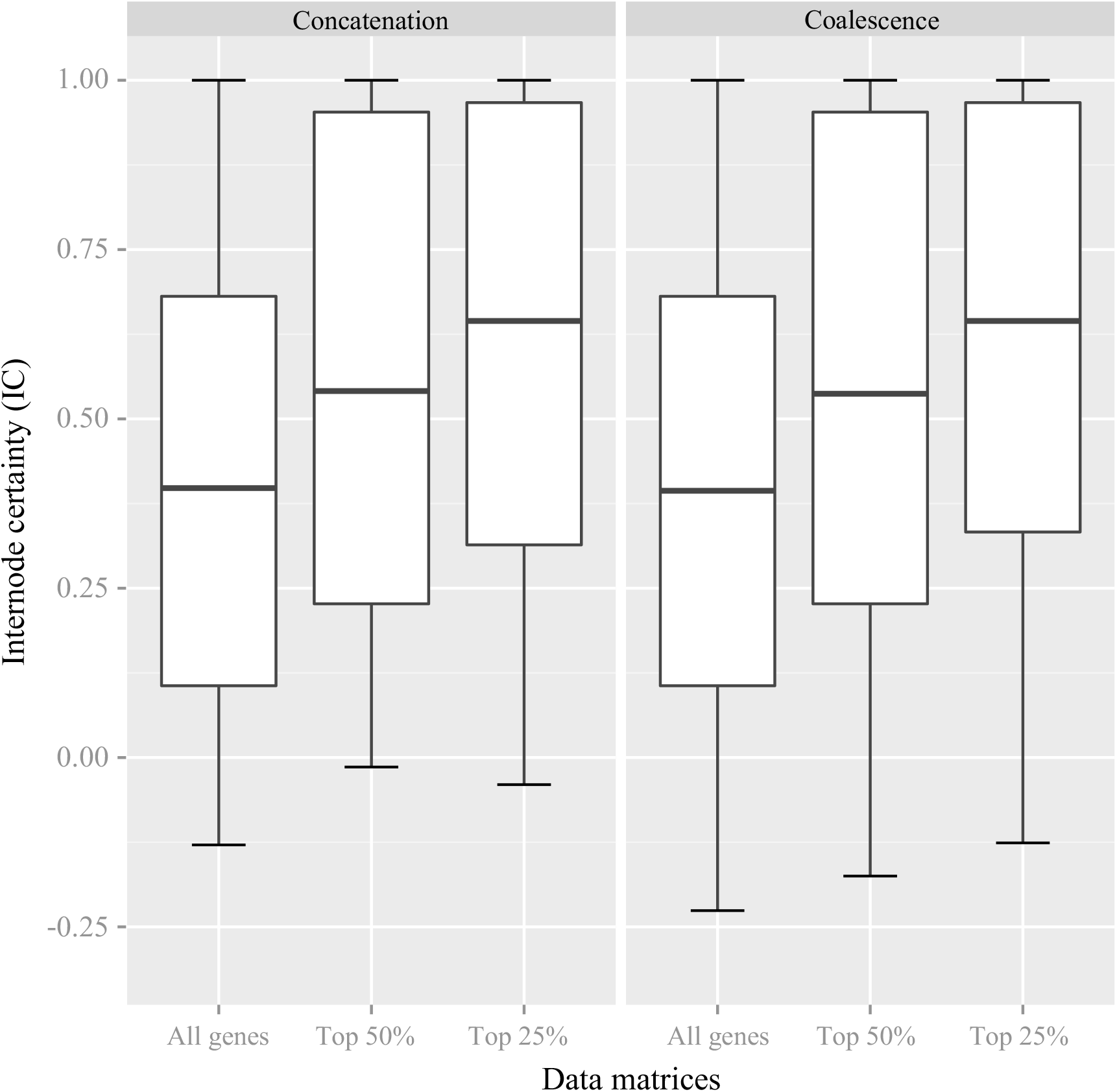
Comparison of the distributions of internode certainty (IC) values across three data matrices and two approaches. The three data matrices are: “All genes”, all 1,233 genes in the AA data matrix; “Top 50%”, AA data matrix including the 616 genes with the strongest phylogenetic signal; “Top 25%”, AA data matrix including the 305 genes with the strongest phylogenetic signal (see Materials and Methods). For each data matrix, the set of individual ML gene trees are used to calculate (partial) internode certainty (IC) values for all internodes in the concatenation-based ML phylogeny (left panel) and the coalescence-based ASTRAL phylogeny (right panel), respectively. Rectangles in the boxplot denote 1st and 3rd quartiles. Horizontal thick bars represent mean IC values.

In agreement with the IC results, there were fewer topological differences between the concatenation- and coalescence-based phylogenies in the two reduced data matrices relative to the full data matrix (5 topological differences instead of 8). These five remaining conflicting internodes occurred within the CUG-Ser clade and the *Yarrowia* clade (Figures S8 and S9). Specifically, the concatenation phylogeny interfered from top 50% data matrix was topologically identical with that inferred from the complete data matrix, albeit more weakly supported. Furthermore, unlike the coalescence-based phylogeny recovered from the complete data matrix, the coalescence-based phylogeny from the top 50% data matrix recovered the *Nakaseomyces* clade as the sister group to the genus *Saccharomyces* (Figure S8). For the top 25% data matrix, both concatenation and coalescence-based phylogenies supported this sister relationship. However, they consistently supported the paraphyly of the WGD clade (Figure S9), which was not recovered in concatenation and coalescence-based phylogenies inferred from the top 50% data matrix but was observed in previous multigene studies (Kurtzman and Robnett 2013; Mühlhausen and Kollmar 2014).

In summary, 74 / 83 internodes in this 86-taxon phylogeny of the Saccharomycotina yeasts are robust to different approaches (concatenation versus coalescence) and phylogenomic data matrices (AA versus C12), while the remaining 9 internodes are still unresolved or equivocal (Figure S10).

## Conclusion

Twenty years ago, the genome sequence of *S. cerevisiae* (Goffeau *et al.* 1996) ushered yeast biology into the age of genomics. Although we still lack genomic data from most of the known yeast biodiversity, the public availability of dozens of yeast genomes provided us with an opportunity to examine the quality of genomic data presently available for the lineage (Figure 2) and infer the backbone phylogeny of the Saccharomycotina (Figures 3 and 4). With several large-scale efforts to sample yeast biodiversity currently underway, such as the 1002 Yeast Genomes Project focusing on *S. cerevisiae* (http://1002genomes.u-strasbg.fr), the iGenolevures Consortium (http://gryc.inra.fr), and the Y1000+ Project focusing on sequencing the genomes of all known species of the subphylum Saccharomycotina (http://y1000plus.org), the phylogenomic analyses reported in this study provide a robust roadmap for future comparative research across the yeast subphylum, while highlighting clades in need of further scrutiny.

## Acknowledgments

We thank Thomas W. Jeffries, Meredith Blackwell, and the DOE Joint Genome Institute for releasing several genome sequences through MycoCosm prior to their formal publication (Riley *et al.* 2016). Mention of trade names or commercial products in this publication is solely for the purpose of providing specific information and does not imply recommendation or endorsement by the U.S. Department of Agriculture. USDA is an equal opportunity provider and employer. This work was conducted in part using the resources of the Advanced Computing Center for Research and Education (ACCRE) at Vanderbilt University and of the UW-Madison Center for High Throughput Computing. This work was supported by the National Science Foundation (DEB-1442113 to AR; DEB-1442148 to CTH), in part by the DOE Great Lakes Bioenergy Research Center (DOE Office of Science BER DE-FC02-07ER64494), the USDA National Institute of Food and Agriculture (Hatch project 1003258 to CTH), and the National Institutes of Health (NIAID AI105619 to AR). CTH is an Alfred Toepfer Faculty Fellow, supported by the Alexander von Humboldt Foundation. CTH is a Pew Scholar in the Biomedical Sciences, supported by the Pew Charitable Trusts.

## Supplementary Figures

**Figure S1.** Box plots of GC content for the 1,233 nuclear protein-coding BUSCO genes in each of the 96 taxa used in this study. The plots are drawn for all sites (All), first codon positions only (Pos1), second codon positions only (Pos2), and third codon positions only (Pos3), respectively. Note that GC content at Pos3 shows substantial variation among the 96 taxa.

**Figure S2.** Distribution of the best-fitting amino acid substitution models across the amino acid alignments of the 1,233 BUSCO genes used in this study. For each gene, the best-fitting model of amino acid evolution was selected using the Bayesian information criterion (BIC) implemented in ProtTest version 3.4 (Darriba *et al.* 2011). Note that we accounted for among-site rate heterogeneity by only using Gamma (G) rather than Gamma (G) + proportion of invariable sites (I), because the G and I parameters are partially overlapping in phylogenetics.

**Figure S3.** Venn diagrams illustrating the overlaps between subsets (left, Top 25%; right, Top 50%) of 1,233 genes displaying the highest average bootstrap support (ABS) and the highest average relative tree certainty (RTC) in their ML gene trees. Note that the subsets of genes selected by the two measures of phylogenetic signal (ABS and RTC) are not substantially different.

**Figure S4.** Illustration of occupancy for each taxon and gene in our data matrices. Each row corresponds to a taxon and each column corresponds to a BUSCO gene. White coloration denotes a gene’s absence in a given taxon, whereas all other coloration denotes a gene’s presence in a given taxon. The range of gray to black coloration reflects the ratio of gene sequence length for a given taxon to the length of the longest sequence for the same gene across the 96 taxa. Genes exhibiting the highest occupancy are toward the right of the graph, and taxa exhibiting the highest occupancy are toward the top of the graph. Note that gap characters were not considered in our calculations of taxon occupancy.

**Figure S5.** Distribution of gene alignment lengths across the 1,233 BUSCO genes used in this study. The sequence alignment length of each gene was measured after filtering out ambiguously aligned positions; all three codon positions were included in this measurement.

**Figure S6.** The phylogenetic relationships of Saccharomycotina yeasts inferred from the concatenation-based analysis of the C12 data matrix. The ML phylogeny was reconstructed based on the concatenation data set (1,219,798 bp) under an unpartitioned GTR + GAMMA substitution model using RAxML version 8.2.3 (Stamatakis 2014). Branch support values near internodes denote bootstrap support (above) and internode certainty (below) values, respectively. * indicates bootstrap support values greater than or equal to 95%. Thicker branches illustrate conflicts between concatenation-based phylogeny (Figure S6) and coalescence-based phylogeny (Figure S7). Note that branch lengths on the ML tree are given in the ***inset*** at the bottom left.

**Figure S7.** The phylogenetic relationships of Saccharomycotina yeasts inferred from the coalescence-based analysis of the C12 data matrix. The coalescence-based phylogeny estimation was conducted using ASTRAL version 4.7.7 (Mirarab *et al.* 2014). Branch support values near internodes denote bootstrap support (above) and internode certainty (below) values, respectively. * indicates bootstrap support values greater than or equal to 95%. Thicker branches show conflicts between coalescence-based phylogeny (Figure S7) and concatenation-based phylogeny (Figure S6).

**Figure S8.** Conflicts in the phylogenetic relationships of Saccharomycotina yeasts inferred from the concatenation-based (a) and coalescence-based (b) analysis of the 616 genes in the AA data matrix whose bootstrap consensus gene trees had the highest average bootstrap support (top 50%). Branch support values near internodes are indicated as bootstrap support values (internodes without designation have values greater than or equal to 95%). Note that outgroup taxa are not shown.

**Figure S9.** Conflicts in the phylogenetic relationships of Saccharomycotina yeasts inferred from the concatenation-based (a) and coalescence-based (b) analysis of the 308 genes in the AA data matrix whose bootstrap consensus gene trees had the highest average bootstrap support (top 25%). Branch support values near internodes are indicated as bootstrap support values (internodes without designation have values greater than or equal to 95%). Note that outgroup taxa are not shown.

**Figure S10.** Supported and unresolved internodes in phylogeny of Saccharomycotina yeasts. Topology and branch lengths given in the ***inset*** at the bottom left are derived from the concatenation amino acid data matrix (Figure 3). Branch support values near internodes are indicated as bootstrap support value (above) and internode certainty (below), respectively. * indicates bootstrap support values greater than or equal to 95%. Solid lines indicate internodes that are robustly supported by different approaches (concatenation, coalescence) and phylogenomic data matrices (AA, C12). Dashed lines indicate internodes that show conflict or are weakly supported; we consider such internodes to be unresolved.

### Supplementary Tables

**Table S1.** List of all taxa used in this study.

**Table S2.** Summary of assessment of 96 genome assemblies.

**Table S3.** Summary of 1,233-gene, 96-taxon data matrix.

## References

Altschul, S. F., W. Gish, W. Miller, E. W. Myers, and D. J. Lipman, 1990 Basic local alignment search tool. J. Mol. Biol. 215: 403–410.

Borneman, A. R., B. A. Desany, D. Riches, J. P. Affourtit, A. H. Forgan et al., 2012 The genome sequence of the wine yeast VIN7 reveals an allotriploid hybrid genome with *Saccharomyces cerevisiae* and *Saccharomyces kudriavzevii* origins. FEMS Yeast Res.12: 88–96.

Camacho, C., G. Coulouris, V. Avagyan, N. Ma, J. Papadopoulos et al., 2009 BLAST+:architecture and applications. BMC Bioinformatics 10: 421.

Capella-Gutierrez, S., J. M. Silla-Martinez, and T. Gabaldon, 2009 trimAl: a tool for automated alignment trimming in large-scale phylogenetic analyses. Bioinformatics 25:1972–1973.

Darriba, D., G. L. Taboada, R. Doallo, and D. Posada, 2011 ProtTest 3: Fast selection of best-fit models of protein evolution. Bioinformatics 27: 164–165.

Dujon, B., 2010 Yeast evolutionary genomics. Nat. Rev. Genet. 11: 512–524.

Eddy, S. R., 2011 Accelerated Profile HMM Searches. PLoS Comput. Biol. 7: e1002195.

Edwards, S. V., 2009 Is a new and general theory of molecular systematics emerging? Evolution. 63: 1–19.

Felsenstein, J., 1981 Evolutionary trees from DNA sequences: a maximum likelihood approach. J. Mol. Evol. 17: 368–376.

Fitzpatrick, D. A., M. E. Logue, J. E. Stajich, and G. Butler, 2006 A fungal phylogeny based on 42 complete genomes derived from supertree and combined gene analysis. BMC Evol. Biol. 6: 99.

Gibson, B., and G. Liti, 2015 *Saccharomyces pastorianus*: genomic insights inspiring innovation for industry. Yeast 32: 17–27.

Goffeau, A., B. G. Barrell, H. Bussey, R. W. Davis, B. Dujon et al., 1996 Life with 6000 Genes. Science 274: 546–567.

Hall, C., and F. S. Dietrich, 2007 The Reacquisition of Biotin Prototrophy in *Saccharomyces cerevisiae* Involved Horizontal Gene Transfer, Gene Duplication and Gene Clustering. Genetics 177: 2293–2307.

Hittinger, C. T., 2013 Saccharomyces diversity and evolution: a budding model genus.Trends Genet. 29: 309–317.

Hittinger, C. T., A. Rokas, F.-Y. Bai, T. Boekhout, P. Gonçalves et al., 2015 Genomics and the making of yeast biodiversity. Curr. Opin. Genet. Dev. 35: 100–109.

Hittinger, C. T., A. Rokas, and S. B. Carroll, 2004 Parallel inactivation of multiple GAL pathway genes and ecological diversification in yeasts. Proc. Natl. Acad. Sci. USA 101:14144–14149.

Hosner, P. A., B. C. Faircloth, T. C. Glenn, E. L. Braun, and R. T. Kimball, 2016 Avoiding Missing Data Biases in Phylogenomic Inference: An Empirical Study in the Landfowl (Aves: Galliformes). Mol. Biol. Evol. 33: 1110–1125.

HovmÖller, R., L. L. Knowles, and L. S. Kubatko, 2013 Effects of missing data on species tree estimation under the coalescent. Mol. Phylogenet. Evol. 69: 1057–1062.

Huelsenbeck, J. P., J. J. Bull, and C. W. Cunningham, 1996 Combining data in phylogenetic analysis. Trends Ecol. Evol. 11: 152–158.

James, T. Y., F. Kauff, C. L. Schoch, P. B. Matheny, V. Hofstetter et al., 2006 Reconstructing the early evolution of Fungi using a six-gene phylogeny. Nature 443: 818–822.

Katoh, K., and D. M. Standley, 2013 MAFFT multiple sequence alignment software version 7: Improvements in performance and usability. Mol. Biol. Evol. 30: 772–780.

Kobert, K., L. Salichos, A. Rokas, and A. Stamatakis, 2016 Computing the Internode Certainty and Related Measures from Partial Gene Trees. Mol. Biol. Evol. 33: 1606–1617.

Kurtzman, C. P., J. W. Fell, and T. Boekhout, 2011 The Yeasts: A Taxonomic Study. Elsevier Science.

Kurtzman, C. P., and C. J. Robnett, 1994 Orders and Families of Ascosporogenous Yeasts and Yeast-Like Taxa Compared from Ribosomal RNA Sequence Similarities, pp. 249–258 in Ascomycete Systematics: Problems and Perspectives in the Nineties, edited by D. L. Hawksworth. Plenum Press, New York.

Kurtzman, C. P., and C. J. Robnett, 1998 Identification and phylogeny of ascomycetous yeasts from analysis of nuclear large subunit (26S) ribosomal DNA partial sequences. Antonie Van Leeuwenhoek 73: 331–371.

Kurtzman, C. P., and C. J. Robnett, 2007 Multigene phylogenetic analysis of the *Trichomonascus, Wickerhamiella* and *Zygoascus* yeast clades, and the proposal of *Sugiyamaella* gen. nov. and 14 new species combinations. FEMS Yeast Res. 7: 141–151.

Kurtzman, C. P., and C. J. Robnett, 2003 Phylogenetic relationships among yeasts of the “*Saccharomyces* complex” determined from multigene sequence analyses. FEMS Yeast Res. 3: 417–432.

Kurtzman, C. P., and C. J. Robnett, 2013 Relationships among genera of the *Saccharomycotina* (*Ascomycota*) from multigene phylogenetic analysis of type species. FEMS Yeast Res. 13: 23–33.

Kurtzman, C. P., C. J. Robnett, and E. Basehoar-Powers, 2008 Phylogenetic relationships among species of *Pichia, Issatchenkia* and *Williopsis* determined from multigene sequence analysis, and the proposal of *Barnettozyma* gen. nov., *Lindnera* gen. nov. and *Wickerhamomyces* gen. nov. FEMS Yeast Res. 8: 939–954.

Kurtzman, C. P., and M. Suzuki, 2010 Phylogenetic analysis of ascomycete yeasts that form coenzyme Q-9 and the proposal of the new genera *Babjeviella, Meyerozyma, Millerozyma, Priceomyces*, and *Scheffersomyces*. Mycoscience 51: 2–14.

Le, S. Q., and O. Gascuel, 2008 An improved general amino acid replacement matrix. Mol. Biol. Evol. 25: 1307–1320.

Liang, D., X. X. Shen, and P. Zhang, 2013 One Thousand Two Hundred Ninety Nuclear Genes from a Genome-Wide Survey Support Lungfishes as the Sister Group of Tetrapods. Mol. Biol. Evol. 30: 1803–1807.

Libkind, D., C. T. Hittinger, E. Valério, C. Gonçalves, J. Dover et al., 2011 Microbe domestication and the identification of the wild genetic stock of lager-brewing yeast. Proc. Natl. Acad. Sci. USA 108: 14539–14544.

Lin, Z., and W. H. Li, 2011 Expansion of Hexose Transporter Genes Was Associated with the Evolution of Aerobic Fermentation in Yeasts. Mol. Biol. Evol. 28: 131–142.

Liti, G., A.N. N. Ba, M. Blythe, C. A. Müller, A. BergstrOm et al., 2013 High quality de novo sequencing and assembly of the *Saccharomyces arboricolus* genome. BMC Genomics 14: 69.

Liu, Y., J. W. Leigh, H. Brinkmann, M. T. Cushion, N. Rodriguez-Ezpeleta et al., 2009 Phylogenomic analyses support the monophyly of Taphrinomycotina, including *Schizosaccharomyces* fission yeasts. Mol. Biol. Evol. 26: 27–34.

Louis, V. L., L. Despons, A. Friedrich, T. Martin, P. Durrens et al., 2012 *Pichia sorbitophila*, an Interspecies Yeast Hybrid, Reveals Early Steps of Genome Resolution After Polyploidization. G3 (Bethesda). 2: 299–311.

Marcet-Houben, M., and T. Gabaldón, 2015 Beyond the Whole-Genome Duplication:Phylogenetic Evidence for an Ancient Interspecies Hybridization in the Baker's Yeast Lineage. PLoS Biol. 13: e1002220.

Medina, E. M., G. W. Jones, and D. A. Fitzpatrick, 2011 Reconstructing the Fungal Tree of Life Using Phylogenomics and a Preliminary Investigation of the Distribution of Yeast Prion-Like Proteins in the Fungal Kingdom. J. Mol. Evol. 73: 116–133.

Mirarab, S., R. Reaz, M. S. Bayzid, T. Zimmermann, M. S. Swenson et al., 2014 ASTRAL:genome-scale coalescent-based species tree estimation. Bioinformatics 30: i541–i548.

Morel, G., L. Sterck, D. Swennen, M. Marcet-Houben, D. Onesime et al., 2015 Differential gene retention as an evolutionary mechanism to generate biodiversity and adaptation in yeasts. Sci. Rep. 5: 11571.

Mühlhausen, S., and M. Kollmar, 2014 Molecular phylogeny of sequenced Saccharomycetes reveals polyphyly of the alternative yeast codon usage. Genome Biol. Evol. 6: 3222–3237.

Nguyen, N. H., S.-O. Suh, C. J. Marshall, and M. Blackwell, 2006 Morphological and ecological similarities: wood-boring beetles associated with novel xylose-fermenting yeasts, *Spathaspora passalidarum* gen. sp. nov. and *Candida jeffriesii* sp. nov. Mycol. Res. 110: 1232–1241.

Philippe, H., F. Delsuc, H. Brinkmann, and N. Lartillot, 2005 Phylogenomics. Annu. Rev. Ecol. Evol. Syst. 36: 541–562.

Polev, D. E., K. S. Bobrov, E. V Eneyskaya, and A. A. Kulminskaya, 2014 Draft Genome Sequence of *Geotrichum candidum* Strain 3C. Genome Announc. 2: e00956–14.

Priyam, A., B. J. Woodcroft, V. Rai, A. Munagala, I. Moghul et al., 2015 Sequenceserver: a modern graphical user interface for custom BLAST databases. bioRxiv http://biorxiv.org/lookup/doi/10.1101/033142.

Riley, R., S. Haridas, K. H. Wolfe, M. R. Lopes, C. T. Hittinger et al., 2016 Comparative genomics of biotechnologically important yeasts. Proc. Natl. Acad. Sci. USA DOI:10.1073/pnas.1603941113.

Rokas, A., B. L. Williams, N. King, and S. B. Carroll, 2003 Genome-scale approaches to resolving incongruence in molecular phylogenies. Nature 425: 798–804.

Salichos, L., and A. Rokas, 2013 Inferring ancient divergences requires genes with strong phylogenetic signals. Nature 497: 327–331.

Salichos, L., A. Stamatakis, and A. Rokas, 2014 Novel information theory-based measures for quantifying incongruence among phylogenetic trees. Mol. Biol. Evol. 31: 1261–1271.

Scannell, D. R., K. P. Byrne, J. L. Gordon, S. Wong, and K. H. Wolfe, 2006 Multiple rounds of speciation associated with reciprocal gene loss in polyploid yeasts. Nature 440: 341–345.

Scannell, D. R., O. A. Zill, A. Rokas, C. Payen, M. J. Dunham et al., 2011 The Awesome Power of Yeast Evolutionary Genetics: New Genome Sequences and Strain Resources for the *Saccharomyces sensu stricto* Genus. G3 1: 11–25.

Schwarz, G., 1978 Estimating the dimension of a model. Ann. Stat. 6: 461–464.

Seo, T.-K., 2008 Calculating Bootstrap Probabilities of Phylogeny Using Multilocus Sequence Data. Mol. Biol. Evol. 25: 960–971.

Simão, F. A., R. M. Waterhouse, P. Ioannidis, E. V Kriventseva, and E. M. Zdobnov, 2015 BUSCO: assessing genome assembly and annotation completeness with single-copy orthologs. Bioinformatics 31: 3210–3212.

Slot, J. C., and A. Rokas, 2010 Multiple GAL pathway gene clusters evolved independently and by different mechanisms in fungi. Proc. Natl. Acad. Sci. USA 107: 10136–10141.

Song, S., L. Liu, S. V. Edwards, and S. Wu, 2012 Resolving conflict in eutherian mammal phylogeny using phylogenomics and the multispecies coalescent model.Proc. Natl. Acad. Sci USA. 109: 14942–14947.

Stamatakis, A., 2014 RAxML version 8: A tool for phylogenetic analysis and post-analysis of large phylogenies. Bioinformatics 30: 1312–1313.

Stamatakis, A., P. Hoover, and J. Rougemont, 2008 A rapid bootstrap algorithm for the RAxML Web servers. Syst. Biol. 57: 758–771.

Stanke, M., and S. Waack, 2003 Gene prediction with a hidden Markov model and a new intron submodel. Bioinformatics 19 Suppl 2: ii215–ii225.

Sugiyama, J., K. Hosaka, and S.-O. Suh, 2006 Early diverging Ascomycota: phylogenetic divergence and related evolutionary enigmas. Mycologia 98: 996–1005.

Tavaré, S., 1986 Some Probabilistic and Statistical Problems in the Analysis of DNA Sequences, pp. 57–86 in Lectures on mathematics in the life sciences, edited by Miura RM. American Mathematical Society, Providence, RI.

Taylor, J. W., and M. L. Berbee, 2006 Dating divergences in the Fungal Tree of Life: review and new analyses. Mycologia 98: 838–849.

Wang, Y., X. Zhou, D. Yang, and A. Rokas, 2015 A Genome-Scale Investigation of Incongruence in Culicidae Mosquitoes. Genome Biol. Evol. 7: 3463–3471.

Waterhouse, R. M., F. Tegenfeldt, J. Li, E. M. Zdobnov, and E. V. Kriventseva, 2013 OrthoDB: a hierarchical catalog of animal, fungal and bacterial orthologs. Nucleic Acids Res. 41: D358–D365.

Wenger, J. W., K. Schwartz, and G. Sherlock, 2010 Bulk Segregant Analysis by High-Throughput Sequencing Reveals a Novel Xylose Utilization Gene from *Saccharomyces cerevisiae*. PLoS Genet. 6: e1000942.

Whelan, N., K. M. Kocot, L. L. Moroz, and K. M. Halanych, 2015 Error, signal, and the placement of Ctenophora sister to all other animals. Proc. Natl. Acad. Sci. USA 112:5773–5778.

Wickett, N. J., S. Mirarab, N. Nguyen, T. Warnow, E. Carpenter et al., 2014 Phylotranscriptomic analysis of the origin and early diversification of land plants. Proc. Natl. Acad. Sci. USA 111: E4859–E4868.

Wolfe, K. H., D. Armisén, E. Proux-Wera, S. S. Óhéigeartaigh, H. Azam et al., 2015 Clade-and species-specific features of genome evolution in the Saccharomycetaceae. FEMS Yeast Res. 15: fov035.

Wolfe, K. H., and D. C. Shields, 1997 Molecular evidence for an ancient duplication of the entire yeast genome. Nature 387: 708–713.

Xi, Z., L. Liu, and C. C. Davis, 2016 The Impact of Missing Data on Species Tree Estimation. Mol. Biol. Evol. 33: 838–860.

Xi, Z., L. Liu, J. S. Rest, and C. C. Davis, 2014 Coalescent versus Concatenation Methods and the Placement of Amborella as Sister to Water Lilies. Syst. Biol. 63: 919–932.

Yandell, M., and D. Ence, 2012 A beginner's guide to eukaryotic genome annotation. Nat. Rev. Genet. 13: 329–342.

Yang, Z., 1994 Maximum likelihood phylogenetic estimation from DNA sequences with variable rates over sites: Approximate methods. J. Mol. Evol. 39: 306–314.

Yang, Z., 1996 Among-site rate variation and its impact on phylogenetic analyses. Trends Ecol. Evol. 11: 367–372.

